# Gene flow between environmental and enteric bacteria with environmental coselection for metal and antibiotic resistance genes: A case study

**DOI:** 10.1101/2025.09.15.676296

**Authors:** Ana Vieira, Vishnu Prasad Nair R U, Theodoros Giakoumis, Ragunathan Latha, M.S. Muthu, G.Seghal Kiran, Nikolaos Voulvoulis, Joseph Selvin, Shiranee Sriskandan

**Affiliations:** Centre for Bacterial Resistance Biology, Imperial College London, London, SW7 2AZ, UK; Department of Infectious Disease, Imperial College London, London, SW7 2AZ, UK; Department of Microbiology, Pondicherry University, Puducherry 605014, India; Centre for Environmental Policy, Imperial College London, London, SW7 2AZ, UK; Centre for Pollution Research & Policy, Civil and Environmental Engineering, Brunel University London, UB8 3PH, UK; Department of Microbiology, Aarupadai Veedu Medical College and Hospital, Vinayaka Mission’s Research Foundation (DU), Kirumampakkam, Puducherry, 607403 INDIA; Department of Pharmaceutical Engineering and Technology, Indian Institute of Technology (BHU), Varanasi-221005, Uttar Pradesh, India, Department of Food Science and Technology, Pondicherry University, Pondicherry, 605014 INDIA

## Abstract

**Background:** The global threat of antimicrobial resistance (AMR) is driven by multiple factors, including inadequate sanitation and the suboptimal use of antibiotics in healthcare. The environment is recognised as an important and underappreciated reservoir for AMR. Pollutants can exert selective pressure that co-selects for resistant phenotypes, while faecal contamination introduces enteric bacteria capable of acquiring resistance elements that provide a survival advantage.

**Aims:** This study aimed to characterize multidrug-resistant (MDR) bacteria in sediment and water samples collected from an industrial site and an adjacent canal in 2023 and 2024. Selective agar plating was used to isolate MDR bacteria, which were subsequently subjected to whole-genome sequencing and analysed for the presence of antibiotic resistance genes (ARGs), metal resistance genes (MRGs), and plasmid content.

**Results:** Forty-two MDR isolates were retrieved and sequenced over two seasons. Most were Gram negative and most originated from the industrial location; no MDR bacteria were found in a control location. Species included both environmental and Enterobacteriaceae, indicative of faecal contamination by wastewater, Of the 40 bacteria that possessed at least one ARG, 87.5% (35/40) also carried at least one or more MRG. ARG identified on mobile genetic elements included CTXM-15, *bla*-NDM1, *bla*-VIM1, other carbapenemases, and 16sRMTases. MRG encoded resistance to copper, silver, tellurium, arsenic, and mercury, while plasmid assemblies showed evidence of genetic exchange within the environmental ecosystem. Levels of mercury, vanadium, zinc and copper in sediments exceeded international guideline values.

**Conclusion:** Industrial locations harbour a spectrum of pollutants that contribute to the development and spread of AMR. In this case study, heavy metals appear to act as a potentially major selective pressure driving the evolution of multidrug resistance through co-selection. The observed diversity of mobilizable ARGs and MRGs in the environment highlights the potential for such toxic environments to serve as incubators for the emergence of novel, high-risk bacterial clones.

## Introduction

Access to effective antibacterial agents is critical to modern medicine, but even simple procedures could become life-threatening if drug-resistant infections cannot be treated. Globally, the burden of antimicrobial resistance (AMR) is greatest in some of the poorest regions, while these are also the regions with least access to newer antimicrobials, and least able to provide effective preventative measures such as access to clean water and sanitation (1).

India is one of the world’s greatest consumers of antibiotics, and is considered to be a hotspot for emergence of high-risk clones due to a wide diversity of AMR genes found in wastewater (2, 3). The evolution of clinically-important ‘high risk’ clones among *Enterobacteriales* is not well-understood except from studies using human isolates (4, 5). While human use of antibiotics in healthcare and animal husbandry may be major drivers of selection and expansion in clinical settings, the role of the environment in promoting emergence of new bacterial clones should not be under-estimated. (6–8)

Environmental contamination with antimicrobial residues, arising from manufacturing waste (9, 10) incomplete degradation during municipal wastewater treatment (11), and direct disposal of human waste and discharges from hospitals (12) have all been recognised as a drivers of AMR. However, antimicrobial use and residues do not account completely for AMR (6,7). While the potential selective impact of waste from antibacterial manufacturing and healthcare may be readily apparent, pollution from non-antibiotic pharmaceuticals, heavy metals, biocides, detergents, microplastics and are also recognised to play a role (13–16).

Specific pollutants such as biocides and heavy metals potentially contribute to AMR via mechanisms that have a clear genetic basis, via co-selection (14,15). Bacterial resistance to the pollutant may be encoded by a gene that is co-located (linked) within a genetic element with a different gene that encodes AMR; this enables both resistance genes to be transferred to daughter cells or, in the case of mobile genetic elements, to different bacteria. Some resistance genes encoding for example efflux pumps may confer resistance to both a pollutant and antibiotics (cross-resistance), while some pollutants may increase expression of genes conferring AMR, and vice versa (co-regulation). When chemical pollutants are coupled with faecal contamination from wastewater, the risks of AMR are greatly magnified. (17). The reasons for this are manifold. Firstly, resistance elements within environmental bacteria may be transferred to faecal bacteria that acquire greater fitness to survive in a polluted environment; despite being able to survive outside the human host, the latter remain capable of causing human disease. Secondly, if the faecal bacteria themselves are carrying genes encoding resistance to pollutants (including antimicrobials) that confer selective advantage in the polluted environment, these strains will be selected for and expanded in the polluted setting. Furthermore, the genes encoding AMR in faecal bacteria could be potentially transferred into environmental bacteria. Plasmid transfer itself may be augmented in polluted settings, as may expression of genes involved in resistance (18). These hypothetical scenarios are supported by compelling metagenomic and phylogenomic studies describing the evolution of clinically important AMR from environmental sources (18–20).

Industrial estates host diverse manufacturing activities that release pharmaceuticals, acids, heavy metals, and biocides into the environment. The convergence of these waste streams often creates local environmental contamination hotspots, where pollutants frequently co-occur with organic loads from wastewater treatment, especially in regions with intensive pharmaceutical production such as India (7,9,17,). To investigate the link between industry and AMR, we sampled water and sediments from a polluted industrial site in India. We integrated a culture-based approach with subsequent whole-genome sequencing (WGS) of any multidrug resistant bacteria identified, to investigate the impact of the industrial landscape on microbial communities, and the presence of resistance genes. Our approach revealed an unexpected but consistent association between antibiotic resistance genes (ARG) and metal resistance genes (MRG), often involving mobile genetic elements (MGE), with evidence of gene flow between environmental and polluting enteric bacteria.

## Methods

### Site selection and sampling

The study was conducted at an industrial estate that comprised of pharmaceutical API and formulation plants, together with other health, cosmetic, and biotech-related industries, selected as it had been reported to exhibit a high level of AMR (21). To assess the influence of industrial discharges on the prevalence of AMR in the receiving aquatic environment, six sampling sites (CSL1-CSL6) were selected (Figure 1). Effluent and sediment samples were collected from the industrial tank that receives discharges from the pharmaceutical complex common effluent treatment plant as well as an open drain serving the industrial estate (CSL1). Water and sediment samples were taken from an adjacent canal at the point where effluent from the tank overflows into the canal (CSL2) and further south at increasing distances along the canal: 5 km (CSL3); 7.5 km (CSL4); 9 km (CSL5). Finally, a remote lake (CSL6), unaffected by any industrial or domestic discharges was designated as control.

**Figure 1.**
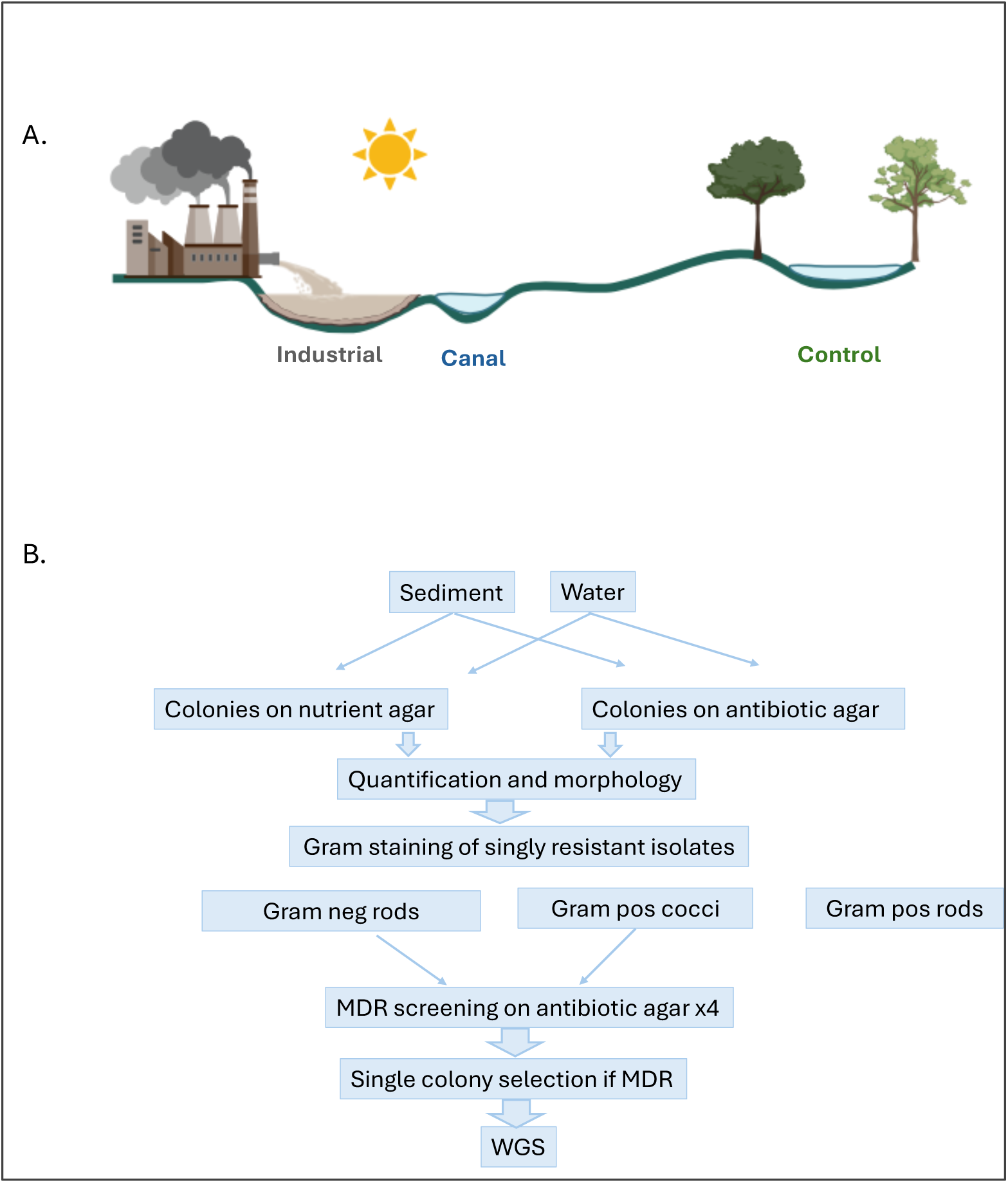
Sampling approach. Sampling sites were from A. Industrial location (CSL1), adjacent canal location (CSL2, CSL3, CSL4, CSL5), and distant control location (CSL6). B. Sediment and water samples (from six sites taken in 2023 and 2024) were plated as indicated, selected on antibiotic agar, and those that were multi-drug resistant (MDR) were selected for whole genome sequencing (WGS). Image created using BioRender.

Samples of water (500ml) and sediment (50ml Falcon tube from a depth of 5 −10 cm) were obtained in triplicate (with replicates obtained 5m apart) in March of 2023 and 2024 at locations that were safe to access. Physicochemical parameters (temperature, pH, oxidation reduction potential, dissolved oxygen, total dissolved solids, salinity, sea water Sigma) of water samples were measured *in situ* using a portable multiparameter probe (Hanna Multiparameter Water Quality Meter, USA).

### Bacteriological analyses

Replicate samples were pooled, then diluted 1:100-1:10,000. 100µl of the 1:100 dilution was plated onto nutrient agar and onto nutrient agar impregnated with one of the following, cefazolin (30µg/ml); cefotaxime (10µg/ml); ciprofloxacin (1µg/ml); ampicillin (25µg/ml); gentamicin (10µg/ml); azithromycin (30µg/ml); and doxycycline(10µg/ml). Plates were incubated at 37^0^C overnight for viable count enumeration.

Colonies growing on antibiotic agar were screened morphologically and up to 5 isolates of each morphology (or 50% if <10 isolates) were selected for Gram staining. Gram negative isolates and Gram positive cocci resistant to one antibiotic were then cross-screened for multi-drug resistance (MDR) by cross-plating onto 4 different antibiotic-impregnated agar plates. Isolates resistant to all 4 agents (cefotaxime, amoxycillin-clavulanate (50 µg/ml), ciprofloxacin, and gentamicin), were designated to be MDR. MDR isolates were then subject to automated antimicrobial susceptibility testing (VITEK 2), and subject to DNA extraction (Himedia HipurA Bacterial Genomic DNA purification kit) prior to whole genome sequencing.

### Genome sequencing and analysis

Whole genome sequencing was carried out using the MiSeq platform with a 300 bp paired-end WGS protocol. The quality of raw reads was assessed using FastQC (22), and reads were trimmed using Trimmomatic (23). The draft genome was de novo assembled using SPAdes v3.13.0 (24). The quality of the assembly was assessed using QUAST v5.0.2 (25), and the assembly was annotated using Prokka v1.14.6 (26) with default settings. Species annotation was performed using two different tools, Kraken2 v2.1.5 (27) and MetaPhlAn2 (28). The potential pathogenicity and ecological niche of the identified bacteria were determined based on a literature search. Genome assemblies were screened for ARG, MRG, and virulence genes (VG) using AMRFinderPLUS v4.0.23 (29) with the “-plus option”. Plasmid analysis was carried out using MOB-suite (30), specifically MOB-recon and MOB-typer tools, with the default options. The identified plasmid contigs were further screened for replicon types using PlasmidFinder Plus (31) and mapped against the PIP-DB database (32) to identify the closest reference plasmids. The completeness of the plasmids was evaluated by mapping the reads against the reference plasmid and visually inspecting them using Proksee (33). Finally, a plasmid and a global phylogenetic tree were constructed to further evaluate the potential transmission of plasmids between strains from different species.

### Chemical analysis of sediments

Metal concentrations in sediment samples (2024 only) from CSL1, CSL2 and CSL6 were measured by Inductively Coupled Plasma Mass Spectrometry (Interfield Laboratories, Kochi, IFL C/QSP G/009 test protocol).

## Results

### Bacteriology

Water and sediment samples from industrial site, canal and control site were used for the study. Samples from the industrial site yielded three orders of magnitude more bacteria than samples from the control site (Supp. Table S1). Among morphologically distinct isolates that were resistant to just one antibiotic, Gram positive bacilli predominated (185/258 isolates from water and 335/486 isolates from sediment). Gram negative bacilli were the next most frequent group among singly-resistant isolates (66/258 isolates from water, and 147/486 from sediments). Gram positive cocci were least frequent (6/258 isolates from water, and 4/486 from sediments). Screening for multidrug resistance (MDR) focused on Gram negative bacteria and Gram positive cocci that were already known to be resistant to at least once antibiotic. In total 12 morphologically distinct isolates from water were MDR, while from sediment, 30 isolates were identified to be MDR. Of these MDR isolates, 32/42 were identified in the industrial location and 10/42 were identified in the canal. No MDR isolates were identified in the control location. Antimicrobial susceptibility testing was undertaken for species where CLSI standards exist; this broadly confirmed the expected phenotypic resistance patterns predicted by the ARGs detected but susceptibility could not be tested for a majority of organisms that are environmental or of unknown clinical significance (Supp Table S2).

### Genome sequencing analysis

Twenty-five different bacterial species were identified including several of clinical relevance and several others that are reported to be opportunistic pathogens to humans (Table 1). Most of the bacteria sequenced were Gram-negative. Known gut commensals included *Klebsiella quasipneumoniae*, *Klebsiella pneumoniae*, *Enterobacter roggenkampii,* and *Enterococcus casseliflavus*. Isolates of *K. pneumoniae* carried up to three plasmids, and *E. roggenkampii,*. carried up to two plasmids (Figure 2, Sup Table S3). Species with reported dual environmental and gut commensalism included *Aeromonas dhakensis*, *Aeromonas simiae*, *Acinetobacter towneri*, and *Acinetobacter pittii*, and *Providencia stuartii* while the remaining strains were considered to be of environmental origin.(Figure 2)

**Figure 2.**
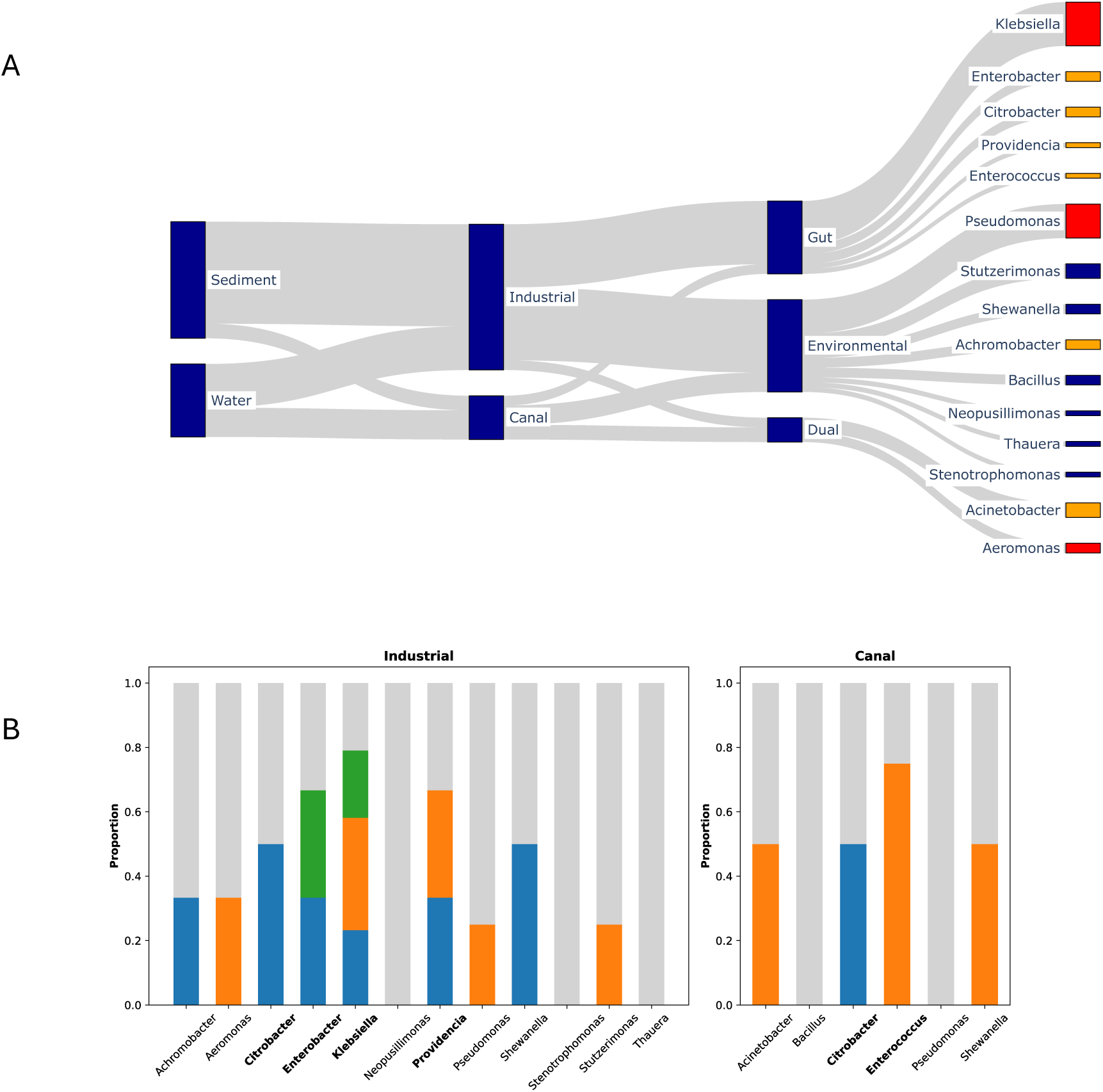
MDR species and plasmid distribution in industrial location and canal. Distribution of cultured MDR bacterial species from sampling locations categorised by usual commensalism (gut, environmental, or dual) **(A).** ‘Dual’ indicates species that have been identified in the enteric flora at least once but are usually considered to be free living. Red tips at the right edge indicate pathogens; orange tips indicate opportunistic pathogens. (**B).** Frequency of different plasmids identified in each species by location. Plasmid types are coloured in a different colour (Blue, ARGs + MRGs; Green, MRGs; Orange, ARGs, Grey – No plasmids) within individual species (enteric species in bold).

**Table 1.** Species identified among MDR isolates. (Upper section 2023; Lower section 2024)

| Strain | Species | Site | Sample | Potential pathogenicity | Reservoir |
| --- | --- | --- | --- | --- | --- |
| MDR1 | *Klebsiella quasipneumoniae* | CSL1 | S | Opportunistic pathogen | Gut |
| MDR2 | *Pseudomonas wenzhouensis* | CSL1 | S | Environmental | Environmental |
| MDR3 | *Pseudomonas wenzhouensis* | CSL1 | S | Environmental | Environmental |
| MDR4 | *Shewanella dokdonensis* | CSL1 | S | Environmental | Environmental |
| MDR5 | *Neopusillimonas aromaticivoran* | CSL1 | S | Environmental | Environmental |
| MDR6 | *Klebsiella quasipneumoniae* | CSL1 | S | Opportunistic pathogen | Gut |
| MDR7 | *Klebsiella pneumoniae* | CSL1 | S | Opportunistic pathogen | Gut |
| MDR8 | *Klebsiella quasipneumoniae* | CSL1 | S | Opportunistic pathogen | Gut |
| MDR9 | *Pseudomonas wenzhouensis* | CSL1 | S | Environmental | Environmental |
| MDR10 | *Pseudomonas wenzhouensis* | CSL1 | S | Environmental | Environmental |
| MDR11 | *Achromobacter spanius* | CSL1 | S | Opportunistic pathogen | Environmental |
| MDR12 | *Achromobacter mucicolens* | CSL1 | S | Opportunistic pathogen | Environmental |
| MDR13 | *Klebsiella pneumoniae* | CSL1 | S | Pathogen | Gut |
| MDR14 | *Klebsiella pneumoniae* | CSL1 | S | Pathogen | Gut |
| MDR15 | *Enterobacter roggenkampii* | CSL1 | S | Opportunistic pathogen | Gut |
| MDR16 | *Pseudomonas wenzhouensis* | CSL1 | S | Environmental | Environmental |
| MDR17 | *Thauera humireducens* | CSL1 | S | Environmental | Environmental |
| MDR18 | *Klebsiella pneumoniae* | CSL1 | W | Pathogen | Gut |
| MDR19 | *Klebsiella quasipneumoniae* | CSL1 | W | Opportunistic pathogen | Gut |
| MDR20 | *Enterobacter roggenkampii* | CSL1 | W | Opportunistic pathogen | Gut |
| MDR21 | *Pseudomonas putida* | CSL1 | W | Opportunistic pathogen | Environmental |
| MDR22 | *Aeromonas dhakensis* | CSL1 | W | Pathogen | Dual |
| MDR23 | *Pseudomonas_khazarica* | CSL1 | W | x | x |
| MDR24 | *Klebsiella pneumoniae* | CSL1 | W | Pathogen | Gut |
| MDR43 | *Aeromonas simiae* | CSL1 | S | Opportunistic pathogen | Dual |
| MDR44 | *Citrobacter sedlakii* | CSL1 | S | Opportunistic pathogen | Gut |
| MDR45 | *Stutzerimonas balearica* | CSL1 | S | Environmental | Environmental |
| MDR46 | *Stenotrophomonas koreensis* | CSL1 | S | Environmental | Environmental |
| MDR47 | *Citrobacter spp.* | CSL2 | S | Opportunistic pathogen | Gut |
| MDR48 | *Bacillus altitudinis* | CSL3 | S | Environmental | Environmental |
| MDR49 | *Bacillus altitudinis* | CSL3 | S | Environmental | Environmental |
| MDR50 | *Acinetobacter junii* | CSL3 | S | x | x |
| MDR51 | *Stutzerimonas stutzeri* | CSL1 | W | Environmental | Environmental |
| MDR52 | *Stutzerimonas balearica* | CSL1 | W | Environmental | Environmental |
| MDR53 | *Providencia stuartii* | CSL1 | W | Opportunistic pathogen | Gut |
| MDR54 | *Franconibacter daqui* | CSL1 | W | Opportunistic pathogen | Environmental |
| MDR55 | *Acinetobacter towneri* | CSL2 | W | Opportunistic pathogen | Dual |
| MDR56 | *Pseudomonas chaetocerotis* | CSL2 | W | Pathogen | Environmental |
| MDR57 | *Shewanella decolorationis* | CSL1 | W | Environmental | Environmental |
| MDR58 | *Enterococcus casseliflavus* | CSL2 | W | Opportunistic pathogen | Gut |
| MDR59 | *Acinetobacter pittii* | CSL2 | W | Opportunistic pathogen | Dual |
| MDR60 | *Acinetobacter pittii* | CSL4 | W | Opportunistic pathogen | Dual |

Industrial site, CSL1; Canal sites. CSL2,3,4. Water, W; Sediment, S. Two Gram-positive bacilli were unexpectedly detected (*Bacillus altitudinis*)

SNP analysis using a reference strain from each respective species showed a high degree of genetic heterogeneity among isolates of the same species. For most of the species, the strains were more than 100 SNPs apart, which suggest the isolates were not related except for isolates of *Citrobacter sedlakii* (13 SNPs apart), *E. roggenkampii* (31 SNPs), *K. pneumoniae* (< 50 SNPs), *K. quasipneumoniae* (24 SNPs), and *Shewanella decolorationis* (1 SNPs). Interestingly, the strains of *C. sedlakii* and *S. decolorationis* were collected in different location, (Industrial site, CSL1 and the canal, CSL2) yet were highly similar, suggesting a cross over of pathogenic organisms from the polluted industrial site to a reservoir of water potentially used recreationally and for agriculture by the local population.

Seventeen different types of plasmid were found across sequenced isolates that were grouped in four potential classes: pathogenicity/virulence (IncFII, IncFIB, IncHI2(A), IncHI1 (A/B)); resistance dissemination (IncX(3/5), IncR, IncA/C2); conjugative plasmids (IncX(3,5), IncP1); and colicin-encoding plasmids (Col3M, ColpVC, Col) (Supp Table S3). Many of these plasmids were in isolates considered to be wholly environmental or capable of free-living in the environment or gut. The predominant plasmid recorded was IncFIB(K) (13/42; occurring in *K. pneumoniae* and *K. quasipneumoniae*, but also in *Citrobacter sedlakii* and *Enterobacter roggenkampii*), followed by IncR (11/42, of which one third were in environmental isolates), IncFIB (pQil) (6/42), IncA/C2 (5/42) and ColpVC (4/42, two of which were in environmental isolates) (Sup Table S3). The most frequent plasmids were distributed among 4 or 5 different species and genera, while the remaining were species- or genus-specific (Figure 3 and Sup Table S3). Species such as *C*. *sedlakii* had a consistent combination of plasmids despite being identified at two different, albeit adjacent, locations.

**Figure 3.**
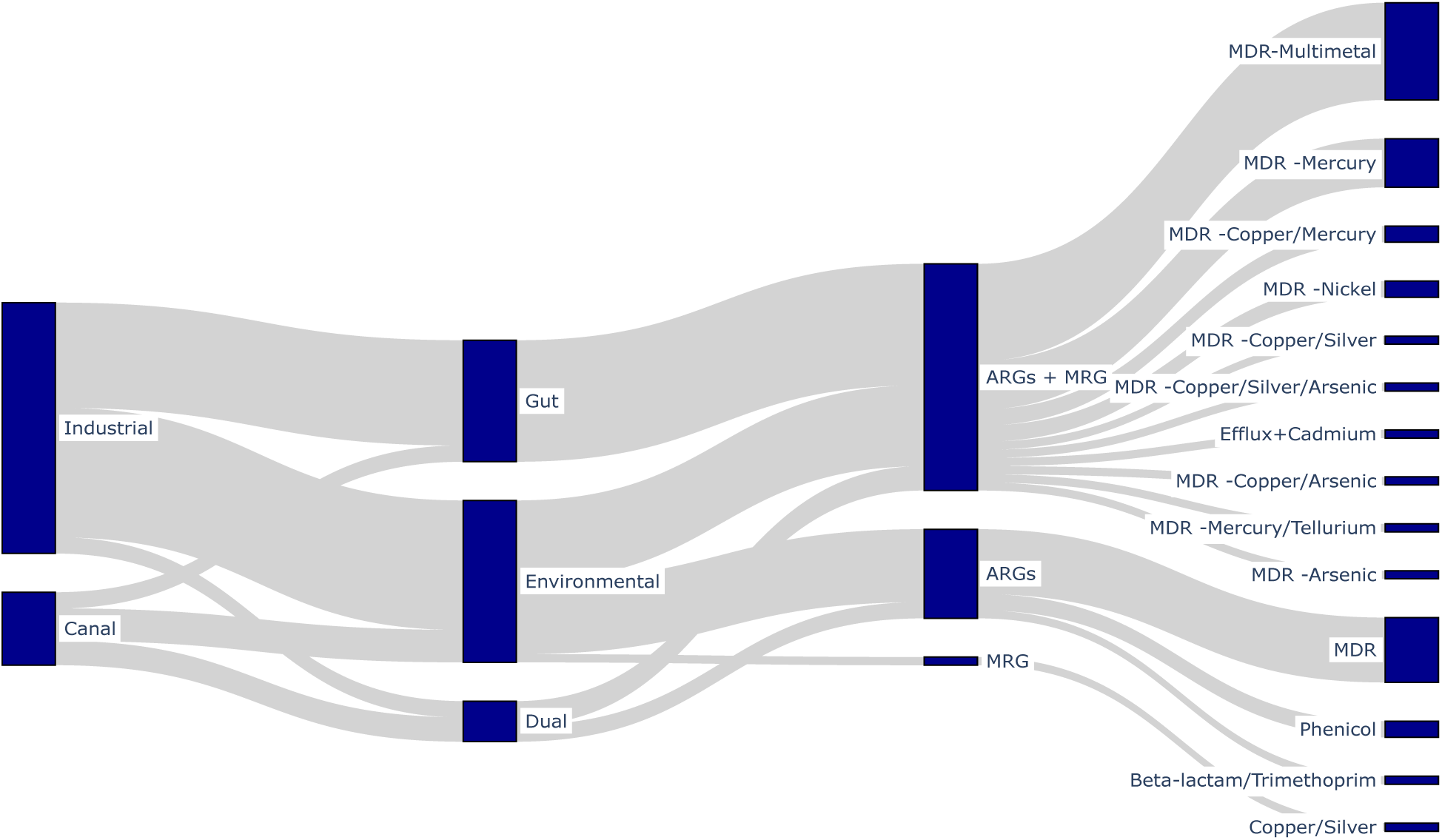
Association between sampling location, species commensalism, and co-location of ARG and MRG.

Genome assemblies were screened for ARG to identify resistance genotypes consistent with MDR status, as well as virulence genes and MRG. Two strains had no ARGs, virulence, or stress genes detected and therefore were removed from the remaining ARG analysis. In total, 130 distinct ARG variants and 40 unique MRG were detected. Remarkably, 87.5% (35/40) of the isolates carried at least one ARG and one MRG (Figure 3), many of which were sited on plasmids some of which are predicted to be conjugative or mobile (Supplementary Figure 1 and Suppl Table S4). Mapping of MRG-carrying plasmid reads to putative plasmid reference sequences showed ∼20% of reads shared between 3 *K. quasipneumoniae*, 2 *K. pneumoniae*, and 2 *E. roggenkampii* isolates to be identical. Approximately 50% of IncF1B reads shared between 3 *K. quasipneumoniae*, and 5 *K. pneumoniae* isolates also appeared identical, although long read sequencing is required to confirm these findings.

Among the ARGs identified, several were clinically important and sited on plasmids occurring in two or more isolates, including *bla*CTXM-15 that encodes for resistance to 3^rd^ generation cephalosporins, the carbapenemase *bla*NDM-1, *AmpC,* and quinolone resistance *qnr* genes. These were associated with MRG in the same isolates (Supp Table S4). Carbapenemase gene *bla*NDM-1 and bleomycin resistance gene *ble* were identified on plasmids in two isolates from the same location, *Acinetobacter towneri* and *Shewanella decolorationis, with A. towneri* possessing 2 additional plasmids and the *bla*OXA-58 carbapenemase. Interestingly both isolates also possessed the same macrolide ARGs msr(E), mph(E), as well as biocide resistance genes *qac*E or *qac*ED1 associated with *sul-*1, that are located on a class 1 integron; *S. decolorationis* additionally possessed 7 mercury resistance genes.

The carbapenemase gene *bla*VIM-1, likely associated with a class 1 integron (indicated by the presence of *qac*E or *qac*ED1 with *sul-*1) (31), was detected in both *Pseudomonas wenzhouensis* and in *K. pneumoniae* from the same location. This strain of *K. pneumoniae* carried a large cargo of ARG and arsenic resistance genes, plus a conjugative IncF plasmid encoding additional resistance genes for copper, silver, arsenic, and antibiotics including CTXM-15.

16sRMTases, a group of enzymes encoding high level aminoglycoside resistance and of key importance in treating carbapenem-resistant Gram negative infections (35), were identified in two *K. pneumoniae* isolates (*rmt*F1), but also in two different isolates of *E. roggenkampii*, where *rmt*B4 was associated with *mcr*9.1. Both isolates possessed additional plasmids loaded with ARGs and genes conferring mercury, copper, silver, tellurium.

### Physico-chemical conditions at sampling locations

The conditions at the industrial location were toxic, with high salinity and poor oxygenation levels. The industrial site water pH ranged from 5.02 (year 1) to 6.82 (year 2), favouring metal ion formation.**(Table 2)**

**Table 2:** Physicochemical properties of water samples.

| **Site** | **Year** | **Temperature**<br>**(°C)** | **pH** | **ORP**<br>**(mVORP)**<br>**Oxidation**<br>**reduction**<br>**potential** | **Dissolved**<br>**Oxygen**<br>**(mg/L)** | **TDS (ppm /**<br>**mg/L)**<br>**Total**<br>**Dissolved**<br>**Solids** | **Salinity**<br>**(psu)** | **Seawater**<br>**Sigma ($\sigma_t$)** |
| --- | --- | --- | --- | --- | --- | --- | --- | --- |
| Industrial<br>CSL1 | 2023 | 25.4 | 5.02 | 18.3 | 2.73 | 14470 | 17.8 | 10.2 |
|  | 2024 | 25.3 | 6.82 | -346.6 | 0.24 | 6639 | 7.61 | 1.8 |
| Canal<br>CSL2 | 2023 | 25.4 | 7.08 | -6.1 | 4.00 | 1,166 | 1.19 | 0.0 |
|  | 2024 | 25.97 | 7.37 | 57 | 1.87 | 611 | 0.61 | 0 |
| Canal<br>CSL 3 | 2023 | 30.3 | 7.70 | 15.4 | 8.18 | 4,745 | 5.28 | 0.0 |
|  | 2024 | 31.32 | 8.44 | 85.1 | 5.02 | 1784 | 1.85 | 0 |
| Canal<br>CSL 4 | 2023 | 33.2 | 7.7 | 2.8 | 6.41 | 5,872 | 6.61 | 0.0 |
|  | 2024 | 33.2 | 8.82 | 92.8 | 6.03 | 20,990 | 26.68 | 14.4 |
| Canal<br>CSL 5 | 2023 | 31.2 | 7.65 | 66.0 | 2.23 | 883. | 0.88 | 0.0 |
|  | 2024 | 35.2 | 8 | 121.4 | 2.66 | 40,790 | 56.67 | 36.1 |
| Control<br>CSL6 | 2023 | 29.2 | 6.65 | 10.5 | 7.25 | 54 | 0.05 | 0.0 |
|  | 2024 | 28.46 | 8.11 | 111.5 | 5.57 | 108 | 0.1 | 0 |

Sediment analysis revealed markedly elevated heavy metal concentrations in both the industrial tank (CSL1) and canal sediments (CSL2) compared to the control site (CSL6), confirming substantial contamination associated with industrial activity. Sediment metal concentrations were obtained in 2024; several exceeded international safety limits in the industrial site (Cu 80.18 mg/kg; Hg 27.89 mg/kg; V 21.15 mg/kg; Zn 277.2 mg/kg) and canal (Cu 18.28 mg/kg; Hg 1.42 mg/kg; V 30.03 mg/kg; Zn 26.63 mg/kg). **(Table 3).** Several metals were present at concentrations more than tenfold higher than the control site, including aluminium, iron, copper, zinc, mercury, and lead (Supplementary Table S5).

**Table 3:** Heavy metals in sediments tested in 2024 exceeding international limits.

| **Heavy metals** | **Industrial**<br>**CSL1mg/kg** | **Canal**<br>**CSL2 mg/kg** | **Control**<br>**CSL6mg/kg** | **International limit**<br>**(sediment)** | **Source** |
| --- | --- | --- | --- | --- | --- |
| Copper | 80.18 | 18.28 | 1.15 | 65 mg/kg | waterquality.gov.au/anz-guidelines |
| Mercury | 27.89 | 1.42 | 0.669 | 0.15mg/kg | waterquality.gov.au/anz-guidelines |
| Vanadium | 21.15 | 30.03 | 2.84 | 7.5mg/kg | echa.europa.eu |
| Zinc | 277.2 | 29.63 | 3.57 | 200 mg/kg | waterquality.gov.au/anz-guidelines |

Limit of quantification (LOQ) was 0.1 mg/kg for all metals except for gold, hafnium, potassium, silicon, sodium, thallium, tungsten where LOQ was 1.0 mg/kg.

## Discussion

Environmental sampling and culture-based analysis were undertaken to determine the burden of multidrug antimicrobial resistance (MDR) in one industrial location that is known to have been polluted. The study identified 42 distinct bacterial isolates of both environmental and faecal origin, that were resistant to three classes of antibiotic; these were identified in the industrial location and adjacent canal, but not in the control location. The MDR isolates not only exhibited an abundance of ARG but also an unexpected abundance of MRG. We posit these genes are in part driven by contamination of the industrial environment with toxic concentrations of heavy metals including mercury, copper, and zinc, and elevated levels of other heavy metals, alongside any antibiotic residues. Importantly ARG and MRG were found on plasmids that are mobilizable, and in both environmental and gut commensal bacteria, pointing to a flow of genetic material between the two bacterial groups.

Environmental bacteria have long been exposed to toxic metals through natural geological processes, driving the evolution of MRGs, that may act via detoxification systems or efflux pumps.(36) These genes, which pre-date the antibiotic era, are frequently located on mobile genetic elements capable of acquiring and mobilising additional resistance determinants. (36) Notable examples include silver (*sil*) and copper (*pco*) operons co-located with multiple ARGs on plasmids, and mercury-resistance transposons commonly linked to integrons carrying ARG payloads. (37)

The co-occurrence of ARGs and MRGs within the same isolates supports the hypothesis that pollution promotes resistance through co-selection mechanisms, as previously demonstrated in environments impacted by mining, wastewater, and industrial effluents (36–39). Furthermore, our findings that near identical combinations of ARGs, MRGs and MGEs exist within different bacterial species in the same environment provides compelling evidence that horizontal transmission of such elements occurs, potentially driven by metal co-selection. Unlike antibiotics, metals can remain in the environment for prolonged periods, exerting long-lasting selective pressures and stabilizing ARGs even in the absence of antibiotics, as well as driving significant shifts in microbial community composition. As such, industrial pollution not only changes microbial ecology but also creates conditions favourable for the persistence and dissemination of resistance (7,14).

Our sediment analysis revealed elevated concentrations of heavy metals in the industrial tank (CSL1) and canal sediments (CSL2), consistent with reports from other pharmaceutical hubs in Pakistan, India and Europe; some were found to be above the maximum permissible limits recommended (10, 40–42). Polluting metals originate from sources such catalysts and other reagents used during the synthesis of pharmaceutical ingredients and the excipients (e.g., stabilizers, fillers, binders, release agents, flavors, colors and coatings), or leaching from manufacturing equipment or pipes (10). Additionally some pharmaceuticals and medical reagents incorporate metals and metalloids by design. These include antimicrobials containing iron, silver and gold; imaging agents using barium, gadolinium, iron, manganese and sodium; lithium for psychotic illness; and platinum-based chemotherapy agents. (43).

The cargo of ARG and MRG within environmental bacteria in our study was associated with class 1 integrons and conjugative plasmids. Furthermore, potentially pathogenic faecal bacteria in the same location had the same, or similar cargo, and an overlapping array of mobile genetic elements. The findings point to complex gene transfer mechanisms involving chromosomal integrons and other mobilizable plasmids, that may allow transfer of ARG and MRG from environmental to pathogenic bacteria, driven by metal co-selection. We propose that contaminated industrial environments may act as genetic ‘nurseries’ or incubators for new high risk AMR clones that are of clinical relevance. The mixing of environmental bacteria with faecal bacteria may, therefore provide an additional hazard for diversification of AMR, underlining the potential value of preventing faecal pollution. However, even where wastewater from industrial sites is treated to remove impurities and reduce ARG, the risks may persist (44). For example, faecal bacteria may enter the environment through pollution from wastewater that may freely drain into the industrial site, as was observed in CSL1. The culture-based approach used in this work provides an unprecedented opportunity to experimentally assess the potential for such genes to be trafficked between donor or recipient environmental and human-pathogenic species, and to establish the mechanisms that might permit this. Understanding the range of potential donor and recipient bacteria in the environmental setting is critical to evaluation of the hazard posed by pollution to AMR.

There are further gaps in our understanding of AMR co-selection and its impact. Despite extensive research on heavy metals in the environment and their potential role in shaping antimicrobial resistance (AMR), there has been little systematic evaluation of the penetration of MRG and co-resistance into clinical and livestock-associated bacterial populations. Environmental AMR studies increasingly rely on metagenomic or PCR-based approaches (17,37), whereas clinical epidemiological research has largely focused on phenotypic testing of isolates with or without genome sequencing (45,46). The contribution of metal-driven co-resistance to veterinary and clinical AMR therefore warrants urgent investigation, particularly given the widespread use of metals such as zinc in livestock feed and their application in medical devices as antibacterial agents.

There were limitations to our study. Samples were collected exclusively during the dry season, limiting our ability to assess potential seasonal effects on the microbiome composition and metal pollution at the industrial site. Moreover, the reliance on a culture-based identification of multidrug resistant bacteria followed by whole genome sequencing limits the analysis to cultivable organisms, overlooking the non-cultivable bacteria that can be detected through metagenomics (47). Despite the efficacy of MOB-Suite to reconstruct plasmid sequences from Illumina short read data (48), ambiguous contigs are assigned to the chromosome, underestimating the number of ARG assigned to plasmids and compromising the completeness of them. To confirm the potential transfer of ARG and MRG between individual bacteria, long read sequencing is required. Finally, although we were able to detect heavy metals in local sediments, quantification was limited to one year, and replicate samples were not tested. Furthermore, total metal content may not represent biologically active ionised metals.

The options to reduce co-selection of AMR by metals in the environment are limited, considering the overlapping pollution pathways across wastewater systems globally. Removing metals from the environment is challenging and costly. Although stricter policies may limit discharges, permissible thresholds still allow the release of metal-contaminated effluents. Moreover, some metal pollution originates from natural geological processes, contributing to background levels of contamination that are difficult to control. Importantly, although the environment may harbour MRG and ARG, these elements are unlikely to transfer into species capable of infecting humans unless environmental bacteria come into contact with enteric (faecal) bacteria. Preventing such faecal contamination represents a more achievable intervention point, through improved sanitation, wastewater treatment, and sewage management, although these measures are unlikely to mitigate transmission within wild ecosystems.

Despite the availability of preventive strategies to curb antimicrobial resistance (AMR), including WASH interventions, vaccination, infection prevention, and antimicrobial stewardship, the burden of AMR, particularly carbapenem resistance, remains alarmingly high in India (45). Our fundings highlight that environmental metal-driven co-selection of AMR poses an additional and underappreciated threat with potential global consequences, underscoring the urgent need to embed One Health–based surveillance programs in regions heavily affected by industrial and faecal pollution.

## Acknowledgements

SS acknowledges the support of the NIHR Biomedical Research Centre awarded to Imperial College London

## Supplementary Data

### Supplementary Figure

**Supplementary Figure S1.**
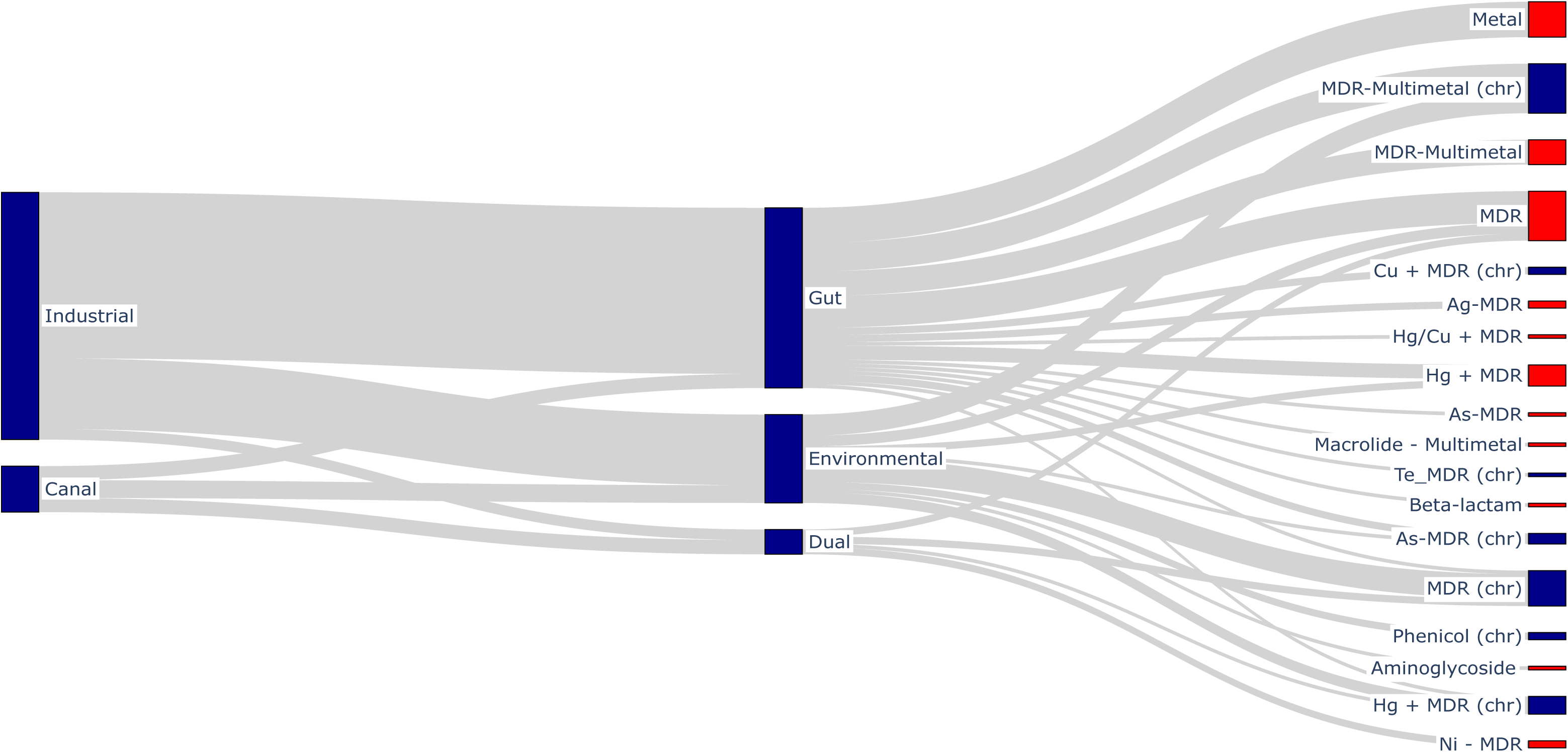
Association between sampling location, species commensalism, and co-location of ARG and MRG. Association with plasmid is indicated by a red tip. Abbreviations Hg, mercury resistance; Ni Nickel resistance; As, arsenic resistance; Te, tellurium resistance; Cu, copper resistance; Ag, silver resistance;

### Supplementary Tables

**Supplementary Table S1.** Bacteriological counts following plating of samples on antibiotic-impregnated agar.

|  | Sediment (per 1 g) |  |  |  |  |  |  |  | Water (per 1ml) |  |  |  |  |  |  |  |
| --- | --- | --- | --- | --- | --- | --- | --- | --- | --- | --- | --- | --- | --- | --- | --- | --- |
| Site | Blank | Cefaz | Cefotax | Cipro | Amp | Gent | Azithro | Doxy | Blank | Cefaz | Cefotax | Cipro | Amp | Gent | Azithro | Doxy |
| **CSL1** | TMTC | TMTC | 250 | TMTC | TMTC | 22 | 146 | 27 | 175 | 130 | 0 | 0 | 0 | 0 | 1 | 0 |
| **CSL2** | 180 | 130 | 10 | 23 | 43 | 0 | 21 | 0 | 80 | 75 | 0 | 9 | 9 | 0 | 0 | 0 |
| **CSL3** | 138 | 16 | 2 | 5 | 29 | 1 | 28 | 0 | 0 | 0 | 0 | 0 | 0 | 0 | 0 | 0 |
| **CSL4** | 3 | 0 | 0 | 0 | 7 | 0 | 0 | 0 | 0 | 0 | 0 | 0 | 0 | 0 | 0 | 0 |
| **CSL5** | 126 | 0 | 0 | 0 | 0 | 0 | 0 | 0 | 0 | 0 | 0 | 0 | 0 | 0 | 0 | 0 |
| **CSL6** | 289 | TMTC | 0 | 0 | 0 | 0 | 0 | 0 | 15 | 251 | 0 | 0 | 0 | 0 | 0 | 0 |

**2024**
|  | Sediment (per 1 gm) |  |  |  |  |  |  |  | water (per 1ml) |  |  |  |  |  |  |  |
| --- | --- | --- | --- | --- | --- | --- | --- | --- | --- | --- | --- | --- | --- | --- | --- | --- |
| Site | Blank | Cefaz | Cefotax | Cipro | Amp | Gent | Azithro | Doxy | Blank | Cefaz | Cefotax | Cipro | Amp | Gent | Azithro | Doxy |
| **CSL1** | TMTC | TMTC | TMTC | TMTC | TMTC | 160 | 250 | 200 | TMTC | TMTC | 75 | 230 | 250 | 124 | 40 | 120 |
| **CSL2** | 240 | 120 | 3 | 0 | 0 | 0 | 3 | 0 | 1 | 12 | 0 | 0 | 0 | 0 | 0 | 0 |
| **CSL3** | TMTC | 200 | 1 | 0 | 0 | 0 | 0 | 0 | 17 | 0 | 0 | 0 | 0 | 0 | 0 | 0 |
| **CSL4** | 20 | 0 | 0 | 2 | 0 | 0 | 0 | 0 | 2 | 0 | 0 | 0 | 0 | 0 | 0 | 0 |
| **CSL5** | 5 | 0 | 0 | 0 | 0 | 0 | 0 | 0 | 0 | 0 | 0 | 0 | 0 | 0 | 0 | 0 |
| **CSL6** | 5 | 0 | 0 | 0 | 0 | 0 | 0 | 0 | 4 | 0 | 0 | 0 | 0 | 0 | 0 | 0 |

Abbreviations Cefaz, Cefazolin; Cefotax, cefotaxime, Cipro, ciprofloxacin; Amp, ampicillin; Gent, gentamicin; Azithro, azithromycin; Doxy, doxycycline
CSL1, Industrial site; CSL2-CSL5, canal; CSL6 control site. Samples (100 µL) were plated out at 1:100 dilution. TMTC, too many to count (>300 cfu) Where TMTC, approximate quantification was estimated using 1:1000 and 1:10,000 dilutions .

**Supplementary Table S2.**
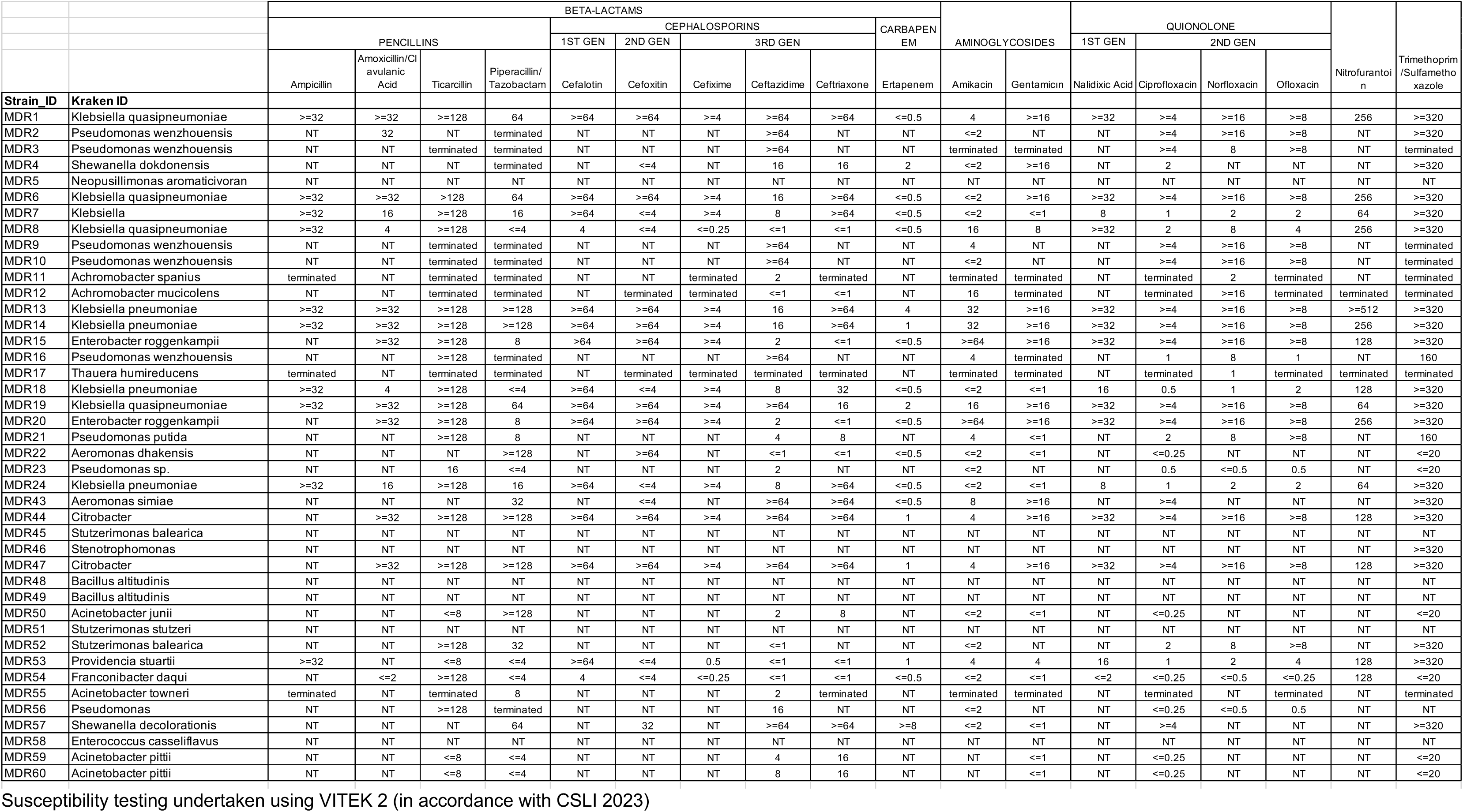
Antimicrobial susceptibility testing of isolates identified to be MDR.

**Supplementary Table S3.**
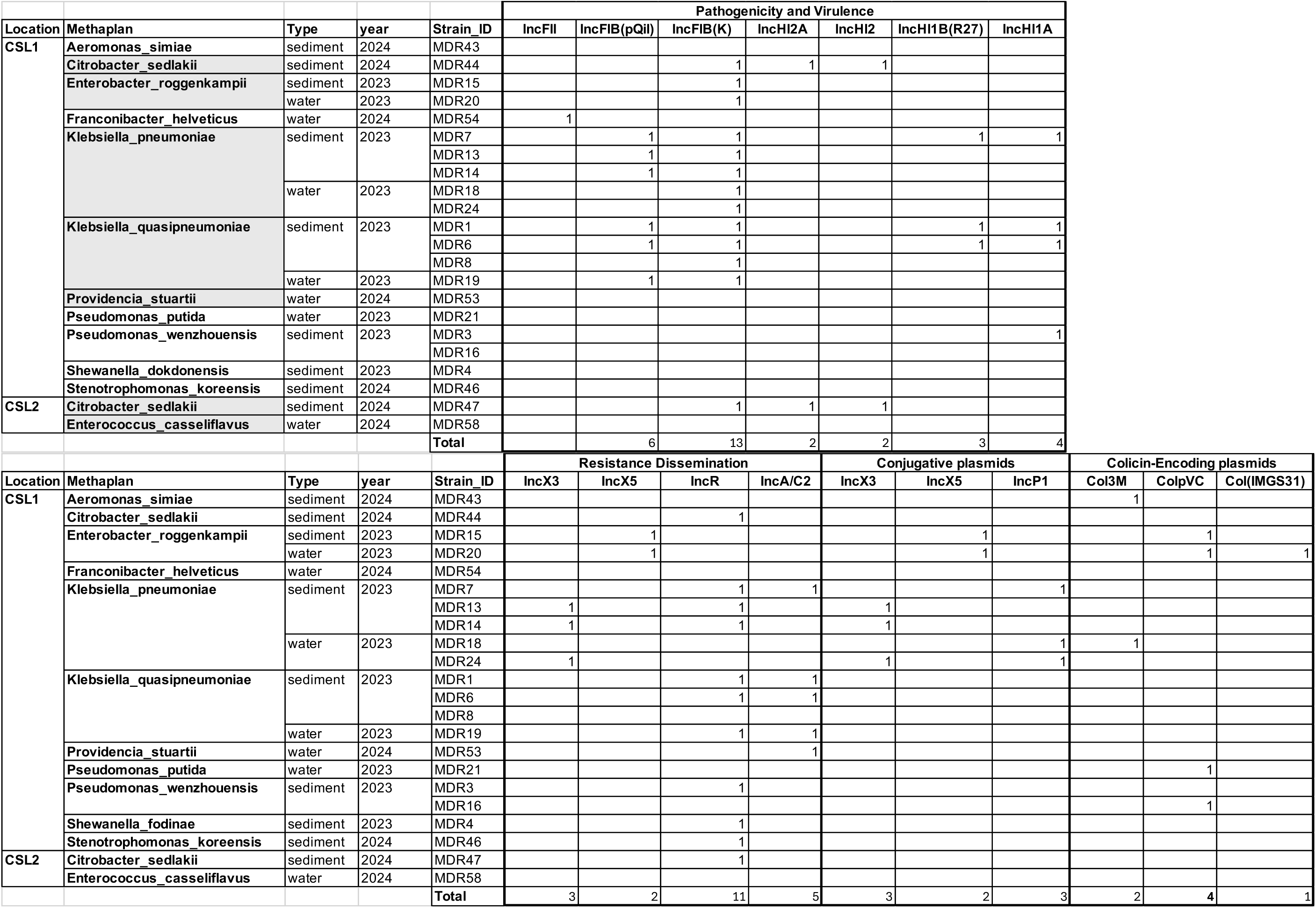
Plasmids identified per bacterial strain.

**Supplementary Table S4.** MDR strains with analysis of plasmids, ARG, and MRG (Excel file)

**Supplementary Table S5.** List of metals quantified in sediment samples in 2024.

| **Heavy metals(mg/kg)** | **Industrial CSL1**<br>**mg/kg** | **Canal CSL2**<br>**mg/kg** | **Control CSL6**<br>**mg/kg** |
| --- | --- | --- | --- |
| Aluminium | 8862 | 10675 | 964.2 |
| Antimony | 0.286 | Not Detected | Not Detected |
| Arsenic | 5.46 | 0.875 | Not Detected |
| Barium | 87.44 | 56 | 3.39 |
| Beryllium | 0.168 | 0.382 | Not Detected |
| Boron | 9.1 | 7.98 | 0.862 |
| Cadmium | 0.12 | Not Detected | Not Detected |
| Caesium | 0.269 | 0.366 | Not Detected |
| Calcium | 2195 | 282.9 | 37.05 |
| Chromium | 48.74 | 28.59 | 9.22 |
| Cobalt | 4.79 | 8.44 | 0.626 |
| **Copper** | **80.18** | **18.28** | **1.15** |
| Gallium | 15.67 | 11.54 | 0.932 |
| Germanium | 0.517 | 0.866 | Not Detected |
| Gold | Not Detected | Not Detected | Not Detected |
| Hafnium | Not Detected | Not Detected | Not Detected |
| Iridium | Not Detected | Not Detected | Not Detected |
| Iron | 6679 | 8093 | 662.6 |
| Lead | 26.28 | 8.38 | 0.175 |
| Lithium | 3.51 | 5.68 | 0.252 |
| Magnesium | 1329 | 2492 | 91.51 |
| Manganese | 82.78 | 191.8 | 19.26 |
| **Mercury** | **27.89** | **1.42** | **0.669** |
| Molybdenum | 4.6 | 1.89 | 0.77 |
| Nickel | 14.82 | 15.65 | 6.04 |
| Niobium | 3.18 | 1.6 | 0.861 |
| Palladium | Not Detected | Not Detected | Not Detected |
| Platinum | Not Detected | Not Detected | Not Detected |
| Potassium | 551.1 | 2044 | 12.21 |
| Rhenium | Not Detected | Not Detected | Not Detected |
| Rhodium | Not Detected | Not Detected | Not Detected |
| Rubidium | 5.5 | 10.57 | 0.137 |
| Ruthenium | Not Detected | Not Detected | Not Detected |
| Selenium | 3.57 | 7.07 | 0.112 |
| Silicon | 707.3 | 623.4 | 210.4 |
| Silver | 3.6 | Not Detected | Not Detected |
| Sodium | 2139 | 85.46 | 38.73 |
| Strontium | 40.18 | 25.42 | 0.95 |
| Tantalum | 0.144 | Not Detected | Not Detected |
| Tellurium | 0.318 | Not Detected | Not Detected |
| Thallium | Not Detected | Not Detected | Not Detected |
| Tin | 1.75 | 0.111 | Not Detected |
| Titanium | 122.1 | 212 | 36.55 |
| Tungsten | 35.06 | 14.38 | 7.91 |
| **Vanadium** | **21.15** | **30.03** | 2.84 |
| **Zinc** | **277.2** | **29.63** | **3.57** |

Bold font indicates concentrations that exceed international recommended limits. Limit of quantification (LOQ) was 0.1 mg/kg for all metals except for gold, hafnium, potassium, silicon, sodium, thallium, tungsten where LOQ was 1.0 mg/kg.

